# PerSEveML: A Web-Based Tool to Identify Persistent Biomarker Structure for Rare Events Using Integrative Machine Learning Approach

**DOI:** 10.1101/2023.10.25.564000

**Authors:** Sreejata Dutta, Dinesh Pal Mudaranthakam, Yanming Li, Mihaela E. Sardiu

## Abstract

Omics datasets often pose a computational challenge due to their high dimensionality, large size, and non-linear structures. Analyzing these datasets becomes especially daunting in the presence of rare events. Machine learning (ML) methods have gained traction for analyzing rare events, yet there remains a limited exploration of bioinformatics tools that integrate ML techniques to comprehend the underlying biology. Expanding upon our previously developed computational framework of an integrative machine learning approach^1^, we introduce PerSEveML, an interactive web-based that uses crowd-sourced intelligence to predict rare events and determine feature selection structures. PerSEveML provides a comprehensive overview of the integrative approach through evaluation metrics that help users understand the contribution of individual ML methods to the prediction process. Additionally, PerSEveML calculates entropy and rank scores, which visually organize input features into a persistent structure of selected, unselected, and fluctuating categories that help researchers uncover meaningful hypotheses regarding the underlying biology. We have evaluated PerSEveML on three diverse biologically complex data sets with extremely rare events from small to large scale and have demonstrated its ability to generate valid hypotheses. PerSEveML is available at https://biostats-shinyr.kumc.edu/PerSEveML/ and https://github.com/sreejatadutta/PerSEveML.

## Introduction

With the continuous expansion of omics and related fields, ML techniques are gaining importance in extracting meaningful insights and advancing our understanding of complex biological systems ^2–5^. Omics data sets encompass large-scale biological data from various disciplines, including genomics, transcriptomics, proteomics, and metabolomics. High-throughput technologies enable researchers to gather a wealth of data from biological samples relatively quickly. Traditional methodologies frequently falter when confronted with the daunting challenges posed by the immense dimensionality, expansive scale, and intricate non-linear structures inherent in omics data ^6^. In this context, ML plays a crucial role in unraveling the intricacies of these vast and complex data sets by deciphering patterns, extracting meaningful insights, and providing actionable intelligence from these multifaceted data repositories.

One of the primary strengths of ML methods is their ability to discover concealed patterns within complex data sets without significant human involvement. This advantage has made ML highly desirable in numerous fields, such as omics data analysis. Additionally, ML strategies are scalable and can process vast amounts of information, making them ideal for high-throughput technologies ^7^. Rather than solely relying on predefined assumptions, ML models learn from the data themselves, which enables them to capture intricate relationships and patterns that traditional analysis methods might overlook. While the ability of ML methods to autonomously uncover hidden patterns is powerful, analyzing rare events in omics data using ML can present several complications ^3^. Most ML algorithms perform poorly when faced with imbalanced data, as they may focus predominantly on the majority class and fail to learn the patterns of rare events, also known as the minority class. ML models typically require sufficient data to learn meaningful patterns. However, in omics data sets, rare events such as specific mutations, rare variants, or low-abundance molecules may be heavily outnumbered by common events ^8^.

Rare events in cancer-genomic studies refer to occurrences of infrequent disease outcomes compared to the controls. For instance, onsite of rare cancers like gallbladder cancer and hairy cell leukemia. In quantitative trait studies, rare events could refer to the expression status of rarely expressed genes or low-abundance proteins ^9,10^. Rarely expressed genes and rare alternative splicing transcripts can provide insights into unique biological processes ^11,12^, low-abundance proteins may indicate specialized biological functions ^13,14^, and rare post-translational modifications (PTMs) can play critical roles in cellular signaling pathways ^15,16^. Analyzing rare events in omics data using ML methods can present several complications due to the inherent challenges posed by the scarcity of these events.

Past research suggested various approaches to deal with the problem of class imbalance, which include data-level, algorithm-based, and hybrid methods ^17,18^. The data-level approaches focus on under-sampling the majority class or over-sampling the minority class before training the data. Thus, this can be considered an extra step in the data preprocessing stage. Some of the well-known sampling techniques include ADASYN ^19^ and SMOTE ^20^. On the other hand, algorithm-based techniques utilize the robustness of some algorithms to deal with class imbalance. Algorithm-based technique includes the concept of cost-sensitive frameworks where a higher penalty for misclassifying is employed on a minority class to improve the classification performance. Another algorithm-based technique includes optimizing the hyperparameters tuning using cross-validation. Cross-validation is a re-sampling procedure to evaluate the performance of an ML model by training several models on subsets of the training set while evaluating them on previously unseen subsets of data ^21,22^. Another method of dealing with class imbalance is the hybrid method. The hybrid methods involve combining data-level methods and algorithm-based approaches. Some prominent hybrid methods include SMOTEBoost ^23^ and RUSBoost ^24^. Often ML practitioners like to use algorithm-based approaches since these approaches are easy to adopt and bring systematic improvement in ML performances without any additional steps. However, it also significantly increases the time to train ML models systematically.

Many analytical tools have been developed in the past decade, specifically designed for high-dimensional omics data. These tools are simple to implement and can present results in a comprehensible format ^25–29^. One such tool is HTPmod ^25^, a web-based shiny application introduced in 2018. HTPmod offers many ML methods and visualization options for high-dimensional data sets. Recently in 2021, another tool called multiSLIDe ^26^ was introduced. This web-based tool visualizes interconnected features between omics data sets for biologists to understand the underlying biology. Another tool to enhance enrichment analysis was developed in 2023 called the Enrichr-KG ^28^. Enrichkr-KG is a web application for gene enrichment analysis and visualization using knowledge graphs. These analytical tools are great research resources due to their easy implementation and require less user input. However, there is still a need for ML tools that address computational challenges related to rare events and visualization techniques that can help researchers formulate meaningful hypotheses.

Analyzing rare events requires careful consideration of data preprocessing, algorithm selection, and validation approaches to ensure robust and meaningful results. However, capturing the essence of rare events can be computationally challenging since limited information is available. To address this challenge, we created PerSEveML, an interactive tool that enables users to predict rare events and visualize how input features contribute to the predictions. The working principle of PerSEveML can be visually depicted through a graphical abstract illustrated in Fig 1.

**Figure 1:**
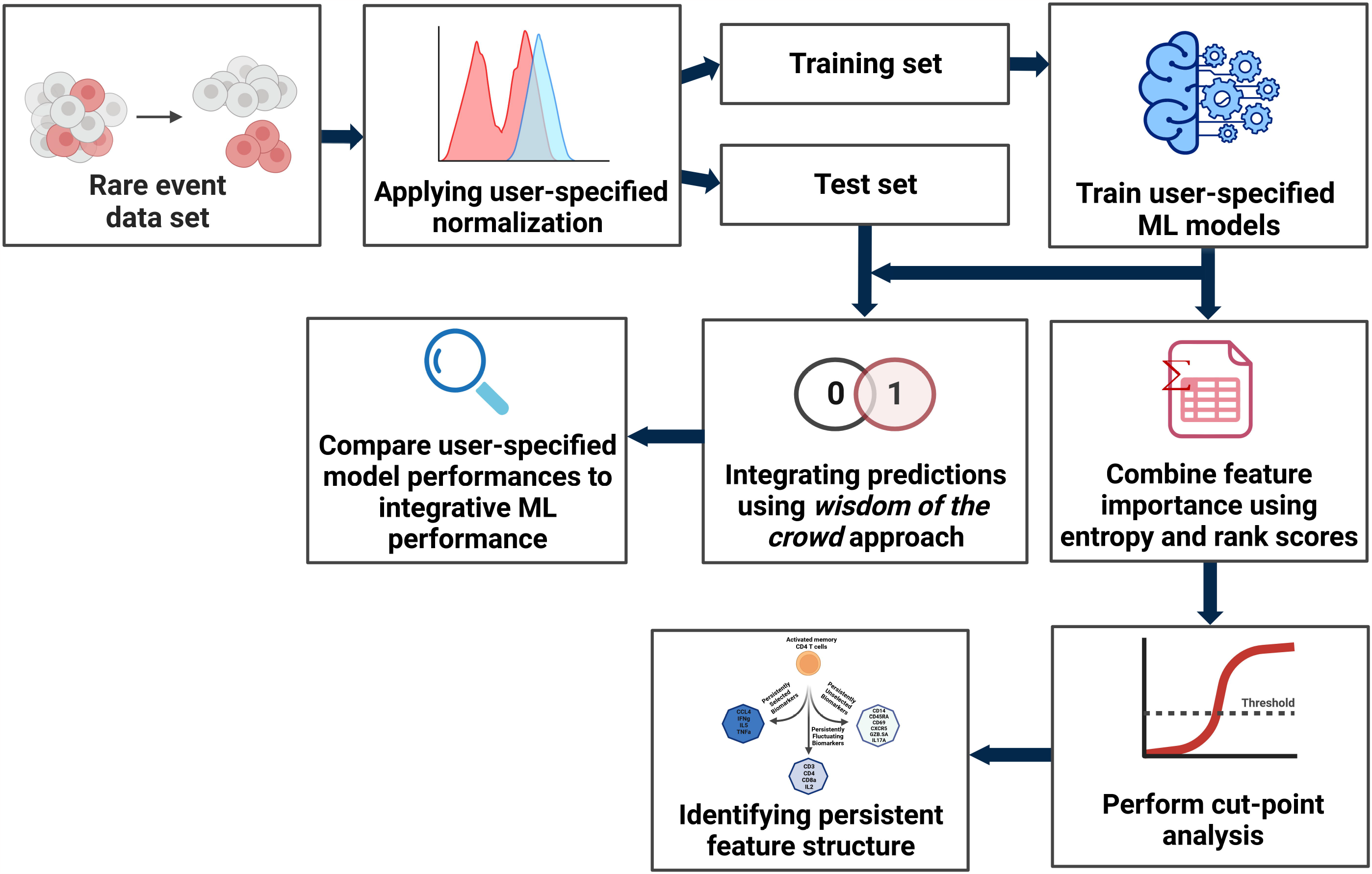
Graphical abstract for PerSEveML framework

PerSEveML is designed to handle common challenges when analyzing omics data sets. Users can choose from twelve ML methods for small to large data sets. Normalization techniques are used to handle non-linear data structures before training ML models. To ensure a wide application of this tool, we have incorporated six different normalization techniques into the interface. The motivation for using six different normalization types is to improve ML model training. For instance, log transformation is often used for skewed data, while standardization is common for feature scaling. TopS, a normalization based on topological scoring, effectively accentuates extreme data points for omics data with rare events ^30,31^. To comprehend the effect of normalization, ML practitioners often rely on data visualization tools such as boxplots. Therefore, we have incorporated box plots into the PerSEveML interface to visualize post-normalization data distribution.

PerSEveML is a versatile tool that can be utilized for various classification problems. What sets it apart is its ability to handle rare events through an integrated ML approach. While other ML toolkits, such as SuperLearner ^27^ and HTPmod ^25^, have attempted to tackle classification problems using multiple ML algorithms, these methods depend on one best-performing model for feature selection, even though more than one model performs comparably. Our goal in adopting an integrative approach was to capitalize on the learning abilities of all top-performing models. Each ML algorithm is influenced by various factors, including cost functions, sampling models, and hyperparameters, which means that different models may identify distinct features that contribute to predicting rare events. Additionally, certain models, such as decision trees, exhibit high variability with low biases.

On the other hand, models like logistic regression or linear discriminant analysis (LDA) display higher biases but lower variances. Since biases and variances can significantly impact an ML model, causing it to be underfitted or overfitted, it is crucial to understand that every ML model used for feature selection or prediction needs to be adequately trained. This ensures that the model gains sufficient exposure to the data to learn all hidden patterns. Furthermore, PerSEveML has been specifically developed to handle complex biological data, such as large protein complex networks with multiple modules and shared subunits. Hence, it is essential to acknowledge that every feature may hold some biological significance.

PerSEveML allows users to assess and compare the performance integrative ML approach with individually selected models using evaluation matrices. The calculation for entropy and rank scores are also available for the user to download and use for further analysis. The app has been designed to address the severe class imbalance issue while ensuring that the ML models are sufficiently trained to prevent underfitting and overfitting. Hence, PerSEveML utilizes algorithm-based cross-validation to deal with class imbalances across the two response categories (binary classification). The users can specify the number of fold *k* for the cross-validation. The integrative approach utilized in this tool has been adapted from 1, where the authors used entropy and rank scores across different ML methods to predict rare events and construct a persistent feature structure, also called persistent biomarker structure.

The PerSEveML interface allows users to visualize the correlation between input features as well as the persistent feature structure created using the integrative ML approach. Our earlier research ^1^, introduced the use of cutpoint analysis to amalgamate feature importance derived from diverse ML methods utilizing entropy and rank score. Thus, the persistent feature structure calculation heavily depends on the cut-off point analysis. The cut-off point can be defined as the percentage of features that users hypothesize to encapsulate the maximum information related to the rare event of interest. Thus, in PerSEveML, we have incorporated the option for users to change the cut-off. This feature allows the users to select the most optimal cut-off point that works for their data set. Following the structure proposed in 1, the proposed feature structure is segregated into persistently selected, fluctuating, and unselected categories. These three categories can be used to select important features from the selected categories or generate a hypothesis using the fluctuating category to understand the association of a weak signaling feature with the rare event being studied.

We highlight the capabilities of PerSEveML by presenting three examples that utilize multi-omics data sets. Each of these data sets has varying sizes and rarity. The first data set is from a study of polychromatic flow cytometry on the rare population of human hematopoietic stem cells (HSCs) ^32^. The cells are derived from human bone marrow cells from a single healthy donor. This data set has 44,140 data points and utilizes expression levels from thirteen (13) surface protein biomarkers to determine the presence or absence of HSC. The second data set is from a high-dimensional flow cytometry and mass cytometry (CyTOF) study on a rare population of activated (cytokine-producing) memory CD4 T cells ^33^. The cells in this data set are derived from human peripheral blood cells exposed to influenza antigens. To determine the presence and absence of T cells, this data set utilizes expression levels from fourteen (14) biomarkers and has 396,460 data points. The third set of data evaluated on PerSEveML consists of proteomics data from 34 focusing on SIN3/HDAC complexes. In this data set, the bait proteins are the features, and the prey proteins are listed in rows. The significance of bait proteins in the complex prediction of SIN3/HDAC complexes has been assessed through protein expression analysis and profiling of the interaction networks of SIN3/HDAC subunits. In summary, we demonstrated the capabilities of PerSEveML as a web tool that simplifies omics data analysis of different sizes, particularly with rare events and enhances the understanding of biological systems.

## Results

### Overview of the application

#### Data preprocessing

PerSEveML is implemented through the R Shiny framework. PerSEveML utilizes the computational power of R while chiming in the in-built parallel computation and user-friendly interface from the Shiny framework. PerSEveML supports multiple file formats including .*RDS* which is generally preferred when dealing with bulky data sets. PerSEveML enables users to select the percentages for training and test sets as part of data preprocessing. PerSEveML supports four common data normalization techniques for omics, including a recently developed TopS normalization ^30^ for all omics data sets with rare events, hyperbolic arcsine with a selectable cofactor for cytometry data sets, log transformation with a flexible constant for skewed features, and percentage row normalization for proteomics data sets. Other normalization options include min-max ^35^, standard scaling or standardization ^36^, and regular arcsine transformation, which can be effortlessly applied across all biological data sets. Users can choose not to use normalization methods if data is already normalized, or normalization is inappropriate for the data set.

Data normalization is an essential part of data preprocessing that allows for optimal training and efficient hyperparameter tuning of ML models. The interface provided by PerSEveML enables users to download and view the normalized data for individual features. Before training ML models, researchers often analyze correlations and heatmaps to check for multicollinearity. PerSEveML simplifies this process by providing correlation plots and the option to download the correlation matrix for further analysis. To ensure that rare event analysis is performed with extra caution, PerSEveML incorporates the concepts proposed in 37. The main foundation of PerSEveML is led by the computational approach proposed in 1. The rationale for choosing the integrative approach was the unique adoption of multiple superior-performing ML methods to understand feature importance rather than relying on one of the best-performing ML methods.

#### Data analysis

To strategize the performance of the integrative approach, PerSEveML can train twelve different ML methods while utilizing the *k*-fold cross-validation as per user choice. The twelve ML methods include tree-based methods such as decision tree, random forest, extreme gradient boosting (XgBoost), and adaptive boosting (AdaBoost); non-tree-based methods include naïve Bayes, linear, non-linear, and polynomial support vector machine (SVM); and linear classifiers such as linear discriminant analysis (LDA), logistic regression, and penalized regression methods such as lasso and ridge ^35,36,38^. Each model is individually tuned to handle the class imbalance problem and evaluated against the test set. PerSEveML allows users to compare the predictive performance of each of these ML models in terms of accuracy, sensitivity, specificity, kappa, and area under the curve for the receiver output characteristics, also known as ROC-AUC or AUC. In addition, PerSEveML calculates the evaluation metrics from an integrative model that combines the predictive capabilities of all the ML models that had been trained.

#### Deriving the persistent feature (or biomarker) structure

In addition to the evaluation metrics, PerSEveML also calculates feature entropies and ranks across all the selected ML models to construct entropy and rank scores. These scores are then compared using a user-defined cut-off to construct a persistent feature structure. As previously discussed, the cut-off point is user-defined, relying on their knowledge of the percentage of biomarkers containing crucial information about rare events. The feature structure consists of three categories: persistently selected, fluctuating, and unselected. The feature structure serves as a feature selection method, where the user can use the persistently selected categories to represent the features that emerged as important features in most of the ML algorithms, suggesting that the feature provides constructive information on the rare event prediction.

On the other hand, the persistently unselected categories represent those features that provided minimal information regarding the rare event, as indicated by the different ML methods. However, the most interesting category belongs to the group of fluctuating features. These dynamic features are pivotal in predicting rare events for certain methods but do not emerge as significant predictors for others. This variation suggests that the complexity of biological processes within these data sets leads different ML models to capture distinct patterns based on their computational algorithm. Hence, these features provide significant hypotheses for future testing. The workflow of PerSEveML is illustrated in Fig 2.

**Figure 2:**
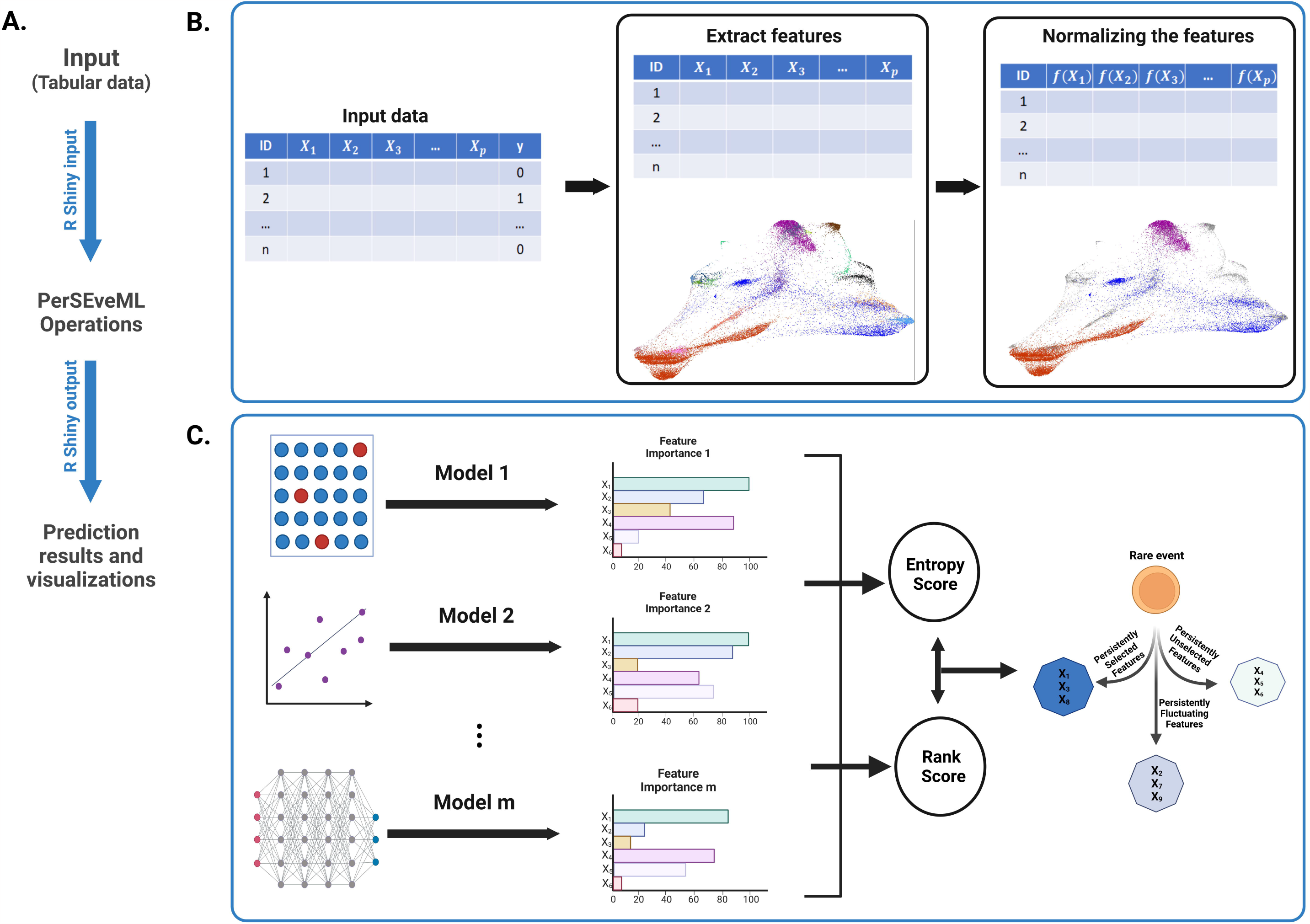
Workflow of PerSEveML

### Predicting rare events and persistent feature structure from PerSEveML

#### Case study 1: Nilsson rare

Nilsson rare represents one large data set with 44, 140 observations and thirteen (13) biomarker expression levels. This data set was introduced in 32 as manually gated data to identify the rare cell population of HSC from human bone marrow. The thirteen surface protein biomarkers included in the study are CD10, CD110, CD11b, CD123, CD19, CD3, CD34, CD38, CD4, CD45, CD45RA, CD49fpur, and CD90bio. 0.8% of the observations, or 358 instances, indicate the presence of rare HSC cells within the 44,140 total observations. Out of these thirteen biomarkers, past studies have confirmed that CD90bio, CD38, and CD45RA play a significant role in identifying HSCs ^39–41^. While surface proteins such as CD11b are expressed on the surface of many leukocytes ^42^, CD45 is mainly expressed in immune cells ^43^.

Using PerSEveML, boxplots revealed that TopS, arcsine transformation with a cofactor of 150, percentage row normalization, and standard scaling all yielded good results when applied to an 80:20% train-test split. However, following the recommendations from 1, our focus primarily centered on TopS, percentage row normalization, and hyperbolic arcsine transformation with a cofactor of 150 to facilitate performance comparison within PerSEveML. Tree-based algorithms, specifically XgBoost, demonstrated commendable performance on this data set, regardless of the normalization method employed. Non-tree-based models followed in performance, with linear models showing the least favorable results. To reach these conclusions, we assessed evaluation metrics such as ROC-AUC, sensitivity, specificity, and kappa.

The performance of the integrative ML approach was highly dependent on the individual models’ performance. For instance, combining a linear model like logistic regression with XgBoost negatively impacted the integrative ML’s performance. However, combining XgBoost with another tree-based model, such as AdaBoost, yielded significantly better results. The feature selection process displayed variations when altering parameters like the training-test split percentage, ‘*k*’ value for *k*-fold crossvalidation, and the cut-off for cut-point analysis. However, CD90bio and CD45RA consistently appeared in the selected feature category, while CD11b and CD45 consistently fell into the unselected category. These findings align with our existing knowledge of HSCs and suggest that the combination of CD90bio, CD34, and CD45RA can reliably identify the presence of HSCs in human bone marrow ^44,45^. The persistent biomarker structure using 80:20 split, TopS normalization, 5-fold cross-validation, and 40% cut-off for cut-point analysis using three ML models (XgBoost, naïve Bayes, and LDA) is presented in Fig 3a.

**Figure 3a:**
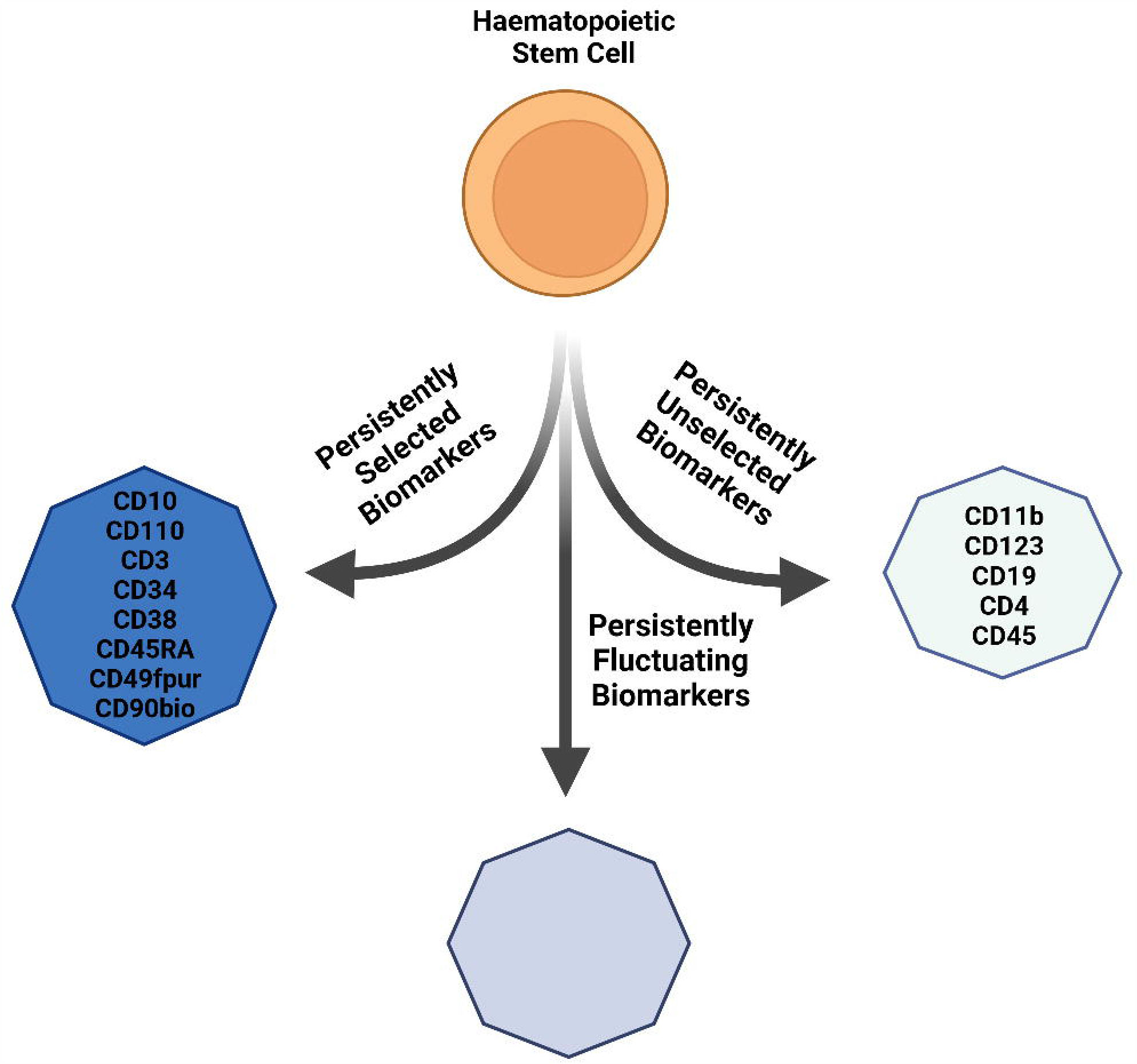
Persistent feature (or biomarker) structure calculated on TopS normalized Nilsson rare data with 80:20 split, 5-fold cross-validation, and 40% cut-off for cut-point analysis using three ML models: XgBoost, naïve Bayes, and LDA

**Figure 3b:**
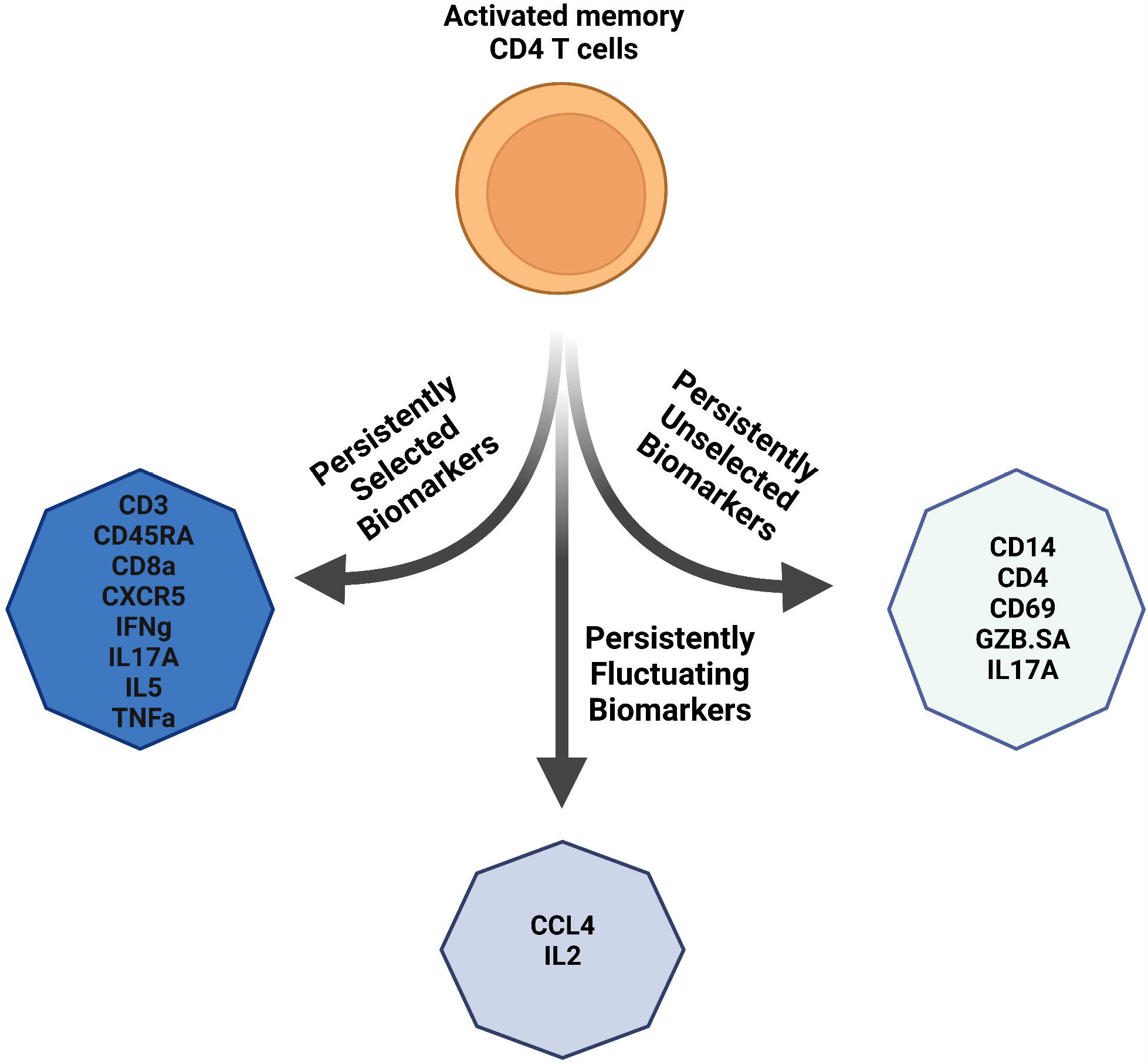
Persistent feature (or biomarker) structure found with Mosmann rare data with percentage row normalization, 80:20 split, 5-fold cross-validation, and 40% cut-off for cut-point analysis using XgBoost and decision tree as ML models

**Figure 3c:**
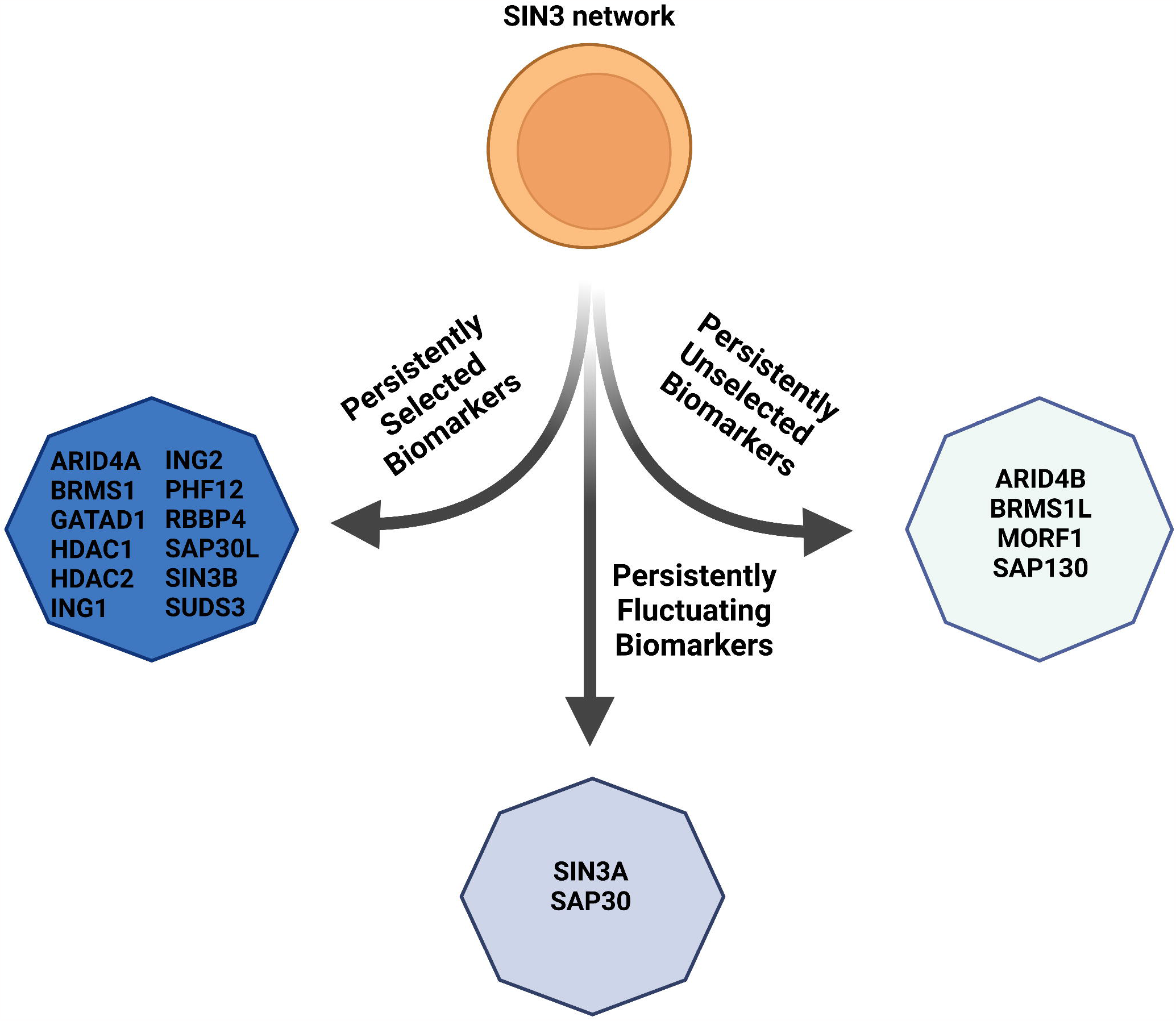
Persistent feature structure illustrated using TopS normalization on SIN3 protein network data, while implementing XgBoost and naïve Bayes as the preferred ML method, and setting a 4-fold cross-validation with cut-point of 30%.

#### Case study 2: Mosmann rare

This extensive data set represents a vast flow cytometry data set comprising 396,460 observations derived from a manually gated data set focusing on a rare population of activated (cytokine-producing) memory CD4 T cells. The data set encompasses fourteen (14) distinct biomarker expression levels. Among these biomarkers, seven (7) pertain to surface proteins, including CCL4, CD14, CD3, CD4, CD45RA, CD69, and CD8a, while the remaining seven (7) are signaling biomarkers, namely CXCR5, GZB.SA, IFNg, IL17A, IL2, IL5, and TNFa.

Hundred and nine (109) cells from the Mosmann rare data set detected the presence of memory-activated CD4 T cells, highlighting an extreme class imbalance with a rarity of 0.03%. Prior research has underscored the crucial role of signaling biomarkers in identifying rare events ^46–48^, with biomarkers such as CD69 proving invaluable in identifying T lymphocytes and natural killer (NK) cells ^49^.

The application of PerSEveML to the Mosmann rare data set revealed that five (5) of the six (6) normalization techniques yielded satisfactory results, with the sole exception being the log transformation, which exhibited relatively underwhelming performance. Similar to the Nilsson rare data set, our focus centered on TopS, percentage row, and hyperbolic arcsine transformations utilizing a cofactor 150 as a normalization technique. Notably, XgBoost consistently demonstrated superior performance compared to all other models.

In general, tree-based models outperformed non-tree-based or linear models. Throughout our iterations, it became evident that signaling biomarkers exhibited superior predictive capabilities compared to surface protein biomarkers. Signaling biomarkers such as IFNg and IL5 consistently stood out as members of the selected category, while CD69 and GZB.SA often found themselves in the unselected category. This underscores the robust performance of PerSEveML in elucidating the underlying biology, as previously suggested by past researchers. The persistent biomarker structure using 80:20 split, percentage row normalization, 5-fold cross-validation, and 40% cut-off for cut-point analysis on the Mosmann data set using two ML methods (XgBoost and decision tree) is presented in Fig 3b.

#### Case study 3: SIN3/HDAC proteomics network

Our latest analysis examined SIN3/HDAC data by using bait proteins as features and listing prey proteins in rows. The data set contains 18 bait and 476 prey proteins, representing their interactions. In this case, rare events refer to prey proteins that are subunits of the SIN3/HDAC complex, which comprise 5.8% of the data. We evaluated the importance of bait proteins in predicting SIN3/HDAC complexes by analyzing protein abundance in interaction networks of SIN3/HDAC complex using data from 34. The task of predicting protein complexes with ML is a complex and challenging one in the fields of bioinformatics and computational biology. Protein complexes are important for various cellular processes, and knowing their composition can provide valuable insights into the functioning of biological systems. However, despite numerous attempts, identifying which human proteins exist in protein complexes and how they are organized on a proteome-wide scale remains challenging ^50^. Recently, ML approaches such as deep learning have been recognized for their potential to predict protein complexes from protein abundances^50^. SIN3/HDAC contains seven homologous pairs: SAP30/SAP30-LIKE, ING1/ING2 (1-like), BRMS1/BRMS1-LIKE, RBBP4/RBBP7, HDAC1/HDAC2, SIN3A/SIN3B, and ARID4A/ARID4B ^34^.

The authors of 34 showed that proteins in homologous pairs may exist in mutually exclusive pairs. Additionally, there are two distinct forms of SIN3 complexes in S. cerevisiae: RPD3L (SIN3 large) and RPD3S (SIN3 small). Higher eukaryote genes encode proteins similar to components of the SIN3 complex in S. cerevisiae. In humans, there are proteins like HDAC1/HDAC2, SIN3A/SIN3B, and RBBP4/RBBP7 that have similarities to the core SIN3 complex componentsRPD3, SIN3, and UME1 in S. cerevisiae. Additionally, humans have proteins similar to components specific to Rpd3L and Rpd3S. For example, SUDS3/BRMS1/BRMS1L, SAP30/SAP30L, and ING1/ING2 have similarities to RPD3L-specific components SDS3, SAP30, and Pho23, respectively. Within RPD3S, components like Rco1 and Eaf3 have similarities to human PHF12 and MORF4L1, respectively. This organization of the SIN3/HDAC complex highlights its complexity, making it an excellent system for ML analysis.

It can be difficult to differentiate between persistently selected and unselected baits when predicting SIN3/HDAC subunits. We experimented with various ML techniques and normalization methods to overcome this challenge. Our findings reveal that we can effectively distinguish between baits by using TopS alongside XgBoost and naïve Bayes and setting a cut-point of 30% and 4-fold cross-validation (Fig 3c). We could also successfully separate mutually exclusive pairs within our data as baits, as illustrated in Fig 3c. For instance, ARID4B was persistently unselected while ARID4A was selected. Similarly, BRMS1L was persistently unselected while BRMS1 was selected. SIN3A/SIN3B, SIN3A was in the fluctuating group along with SAP30 in the SAP30/SAP30L pair. Although the ING1/ING2 pair is traditionally considered mutually exclusive, our data set includes both proteins in the purifications, explaining their presence in persistently selected features. Based on the criteria mentioned earlier, it was discovered that one of the subunits of the large complex, identified as SAP130, was not selected in the persistent group. This particular subunit could not pull down some subunits that compose the SIN3/HDAC complex, distinguishing it from other baits and placing it in the persistently unselected group. In the case of the small complex, the MORF4L1 bait was separated from the PHF12 bait in the persistently unselected group, as it pulled lower abundance proteins overall compared to PHF12 bait.

Thus, these results show that our ML approach applied to protein abundances can reveal hidden features for protein complex prediction that are not easy to detect without prior knowledge.

### Stability and robustness of PerSEveML

The performance of PerSEveML can be likened to the original work by 1, as the computational framework employed in this study draws significant inspiration from the persistent biomarker structure introduced by 1. Nevertheless, it is essential to note a distinction in our approach, as our study primarily centered on a single normalization technique, as opposed to the two normalization methods advocated by the original authors.

Despite this divergence, our findings regarding normalization techniques, the selection of ML methods, and the persistence of the biomarker structure in PerSEveML mirrored those of the original study. As such, we observed that hyperbolic arcsine transformation and TopS work great for both data sets. In addition, both these data sets work well with tree-based ML algorithms, followed by non-tree-based ML approaches. In contrast, the linear models fail to show superior performance. For instance, in the case of the Nilsson rare data set, where we applied the TopS method with an 80:20 percentage split, 10-fold cross-validation, and a cut-off of 40%, we identified XgBoost, naïve Bayes, LDA, and decision tree as models meeting the criterion of sensitivity and specificity above 0.70. This aligns with the outcomes reported by our original paper ^1^, except for the decision tree showing superior performance.

Furthermore, when we employed PerSEveML with the same criteria and cut-off on the Nilsson rare data set, the persistent biomarker structure categorically separated the thirteen biomarkers into either a selected or unselected category (Figure 3a). Notably, none of the biomarkers originally designated as selected or unselected in the original algorithm switched categories. However, an interesting observation emerged: in contrast to the original algorithm, which had fluctuating biomarkers that transitioned evenly between selected and unselected categories using PerSEveML. Thus, PerSEveML revealed more dynamic shifts in these biomarker categories through these case studies.

On further exploration, we observed no significant alteration in biomarker categorization when we applied hyperbolic arcsine transformation or TopS normalization in the Mosmann data set. The key finding is the importance of signaling biomarkers in predicting CD4-activated T cells.

## Discussion

Our research aimed to bridge the gap between the vast resources of ML methods and their applicability to rare events in complex biological processes. The distribution of these rare events presents a computational challenge as they are generally spread across multiple clusters of non-rare events. Topological methods, such as TopS normalization, can condense information related to rare events into a more manageable format for ML models. An important feature of PerSEveML is the normalization methods designed for rare events in omics data sets. Additionally, it integrates multiple ML methods for prediction and feature selection and offers flexibility regarding the train-test split, cross-validation folds, and cut-point analysis. PerSEveML provides complete access to figures and tables for further analysis or publication.

To demonstrate the robustness of PerSEveML in modeling and visualizing results from a crowd-sourced intelligence ML approach, we used three data sets varying in size, number of biomarkers, and rarity percentage. Each data set had unique complexities due to differences in the distribution of rare events across biomarkers. However, by normalizing the data and utilizing high-performing ML methods, we drew informative conclusions on the predictive properties of biomarkers using the persistent feature structure. PerSEveML’s use of entropy and rank scores allows for less rigid results than feature importance generated by individual models. PerSEveML stands out to SuperLearner ^27^ and HTPmod ^25^, which are similar tools, in the context of a more robust feature selection method since it not only utilizes multiple ML methods to predict but combines the strength of pattern recognition from many ML methods to perform feature selection using the persistent feature structure.

PerSEveML automates data analysis with a point-and-click interface. Users must be cautious in drawing valid conclusions from our application output. For example, not all biological processes can be analyzed using TopS or percentage row. Non-tree-based models can perform better than tree-based models, so users should iteratively check which models are used while calculating persistence feature structure. Results may be invalid if users select poorly performing models and draw conclusions based on persistent structure. Users must select ML methods that work best for their data sets and assess multicollinearity prior to training final models.

Even though PerSEveML was built on our previous work of 1 in the realm of persistent biomarker structures, PerSEveML’s computational framework focuses on faster computation and easy ML implementation in analyzing rare events. For instance, we decided to work extensively with a single normalization method. Even though implementing TopS and percentage row normalization is time-consuming for very large data sets, we decided to include them in the application since these normalization techniques are extremely important when working with omics data sets. In addition, unlike 1, we have not included KNN as a part of PerSEveML since, during testing, we found that KNN takes a significantly longer time to tune parameters, and for neither of our test cases did the algorithm show optimal performance. Another deviation from the original work is related to the two methods of calculating feature importance, one via the inbuilt feature importance method from the *caret* package while the other using the stepwise ROC method, introduced by 1. However, stepwise ROC is extremely computationally intensive. Hence, we decided to drop the option from our application.

As demonstrated through the examples of Nilsson and Mosmann’s rare data sets, we found that PerSEveML could capture all the major findings from past articles. However, there were certain differences observed accounting for the persistent feature structure. The differences are because we adapted the algorithm from 1 rather than applying it to its full-scale implementation. Instead of utilizing two techniques of normalizations and variable importance calculation, we used only one at a time. The rationale was the excessive time consumption when incorporating two normalization techniques. We also observed that stepwise ROC-AUC calculation might not be feasible since it is very computationally intensive. Thus, there might be some subtle differences while the method is applied. However, when combined with various ML methods, users can select different normalization techniques to highlight hidden patterns. Since entropy and rank scores are easily downloadable, more curious users can easily implement the original author’s approach and find the more robust version of the persistent biomarker structure.

In essence, our study contributes to the ongoing exploration of persistent biomarker structures, affirming the robustness of PerSEveML in identifying relevant models and highlighting its ability to detect subtle shifts in biomarker categorization. Furthermore, our method showcases PerSEveML’s ability to analyze intricate data structures. For instance, our method can predict stem cells that belong to multiple clustering groups instead of just one. This is evident from our unsupervised clustering analysis in our previous work ^1^. PerSEveML can also analyze protein complexes consisting of modules with shared subunits and mutually exclusive pairs with complex topological structures. As we continue to refine and expand our understanding of these persistent features, PerSEveML stands as a valuable tool for unraveling the complexities of biomarker-driven phenomena, paving the way for more precise and insightful analyses in the field. Finally, it is worth mentioning that our app can be applied to various domains of network-based research.

## Methods

The integrative approach adopted in this study can be segmented into four different parts. The first one is normalization. We have implemented various normalization techniques in our app, including the hyperbolic arcsine transformation, specifically designed for cytometry data. We applied the hyperbolic arcsine transformation with a cofactor of 150, consistent with previous studies. The application features topological scores suitable for multi-omics data, as demonstrated in previous studies ^30,31^. TopS can be mathematically described by equation (1).

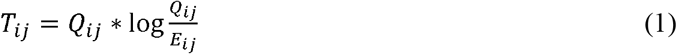

Where, *T*_*ij*_ is the normalized value of *i*^*th*^ biomarker of *j*^*th*^ observation, *Q*_*ij*_ is the expression level of *i*^*th*^ biomarker of *j*^*th*^ observation, and 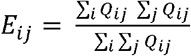

TopS is a topological scoring method that accentuates extreme data points, making it effective for analyzing biologically complex rare event omics data sets. In doing so, TopS helps segregate the rare cell population from the abundant cells and reduces the number of clusters across which the rare cells originate. Past researchers have shown the effect of TopS on different data sets, which can be visually examined through Figure 2 in 1. Since PerSEveML has been developed to analyze more data, we included four more normalization techniques, which include min-max normalization ^35,38^. The mathematical formulation of max-min normalization can be described by equation (2).

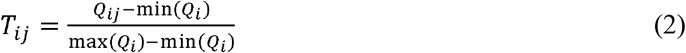

Where, min(*Q*_i_) and max (*Q*_*i*_) are the minimum and maximum values of *i*^*th*^ biomarker across all observations. Min-max normalization usually scales the features between zero (0) and one (1). Min-max normalization is a common preprocessing technique among ML practitioners. PerSEveML also included standardization or the z-score normalization ^35,38^. Equation (3) defines the z-score normalization mathematically.

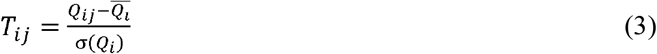

Where 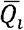 is the mean of the *i*^*th*^ biomarker and σ(*Q*_*i*_) is the standard deviation of *i*^*th*^ biomarker. Standardization transforms individual features into a standard normal distribution with a mean of zero (0) and a standard deviation of one (1). However, this type of normalization fails when the features are skewed. Thus, we also included log transformation, where the users can include a small numeric value beginning from 0.001. Log transformation is very useful when the data is skewed since it can make the transformed features appear like a Gaussian distribution. The sixth type of normalization included in PerSEveML is named a *percentage row*, which can be defined by equation (4).

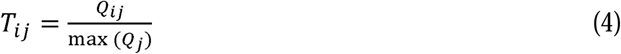

Where, max (*Q*_*j*_) is the maximum value across the *j*^*th*^ observation. This normalization is expected to work similarly to TopS for proteomics data sets.

The next step in data analysis is the selection of the ML methods. PerSEveML is developed for prediction of rare events as well as for feature selection. Thus, all the common classification methods are included in PerSEveML ^35,36,38^. About twelve ML methods incorporated into PerSEveML can be categorized into three classes: tree-based, non-tree-based, and linear classifiers. The tree-based classifiers include decision tree, random forest, XgBoost, and AdaBoost. The non-tree-based methods include naïve Bayes, linear, non-linear, and polynomial SVM. The four linear classifiers included in the model include LDA, logistic regression, and penalized regression methods such as lasso and ridge. Based on the user selection, individual models are tuned to find the optimal hyperparameters using *k*-fold cross-validation. The user can select the number of folds and accept values between 2 and 10.

Once the models are trained, the users can evaluate the performance of the selected models based on evaluation metrics such as sensitivity, specificity, accuracy, kappa, and ROC-AUC. In addition, based on the predictive performance of all the selected models, PerSEveML internally performs a voting classification based on the highest number of predicted classes for individual observations on the test set; the integrative ML approach constructs an integrative prediction. This prediction is also compared to the observed classes on the test set. If all the selected models perform well, the integrative ML shows performance metrics closer to one, while the performance of the integrative ML model lowers when one or more models does not show good performance. Based on these trained models, we then calculate the feature importance using the inbuilt caret package. Instead of using caret package and stepwise ROC to calculate feature importance like 1, we utilized the faster and commonly used method found in literature ^3,35^. PerSEveML calculated the entropies and ranks of individual features based on their importance. Entropy is often associated as a measurement of uncertainty in a system ^1^. To calculate the entropy score across the different ML methods, the adopted mathematical formula is expressed using equation (5).

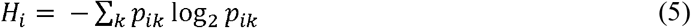

Where, *H*_*i*_ is the entropy score for the *i*^*th*^ biomarker, *p*_*ik*_ is the probability of *i*^*th*^ biomarker in the k^th^ ML model. It should be noted here that to calculate entropy, probabilities need to be calculated. Hence, in case a feature had an importance of zero or negative, the values were replaced by very small constant values such as 10^-12^ and 10^-15^. This is similar to what the original integrative ML approach authors suggested and other studies while conducting their analysis. While the rank scores are also calculated following the formula (equation (6)) from 1.

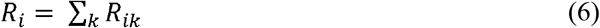

Where *R*_*i*_ is the rank score for the *i*^*th*^ biomarker and *R*_*ik*_ is the rank of *i*^*th*^ biomarker in the k^th^ ML model. The higher the values of rank score and entropy score, the higher the importance of a feature in the predictive model. The next step in data analysis is calculating the persistent feature structure where both these scores are utilized.

Using the cut-off provided by the user, PerSEveML computes the cut-point analysis. Based on the cut-off *c*, PerSEveML selects the top *c%* of features across entropy and rank scores separately. Suppose a feature is in the top c% across both the entropy and rank scores. Then, it is designated as the persistently selected feature since it is a top predictor across many ML methods and two different methods of scoring feature importance. If a feature is not selected in the top c% across both scores, it is considered persistently unselected. Whereas features that show up in the top *c%* in either one of the scoring methods but fail to show up in the other are categorized as persistently fluctuating. The persistent feature structure can not only give users the ability to recognize the most important or least important features but also give users a chance to understand the features that still have weaker associations with the outcome of interest. This might help researchers formulate future hypotheses to understand these weaker associations and the factors associated with them.

## Data availability statement

The mentioned Nilsson and Mosmann data sets are available publicly in FlowRepository (https://flowrepository.org/), repository FR-FCM-ZZPH and can be accessed through 51. While SIN3 data set can be accessed directly from the supplementary materials associated with 34.

The PerSEveML tool is accessible for free at https://biostats-shinyr.kumc.edu/PerSEveML/. For handling larger datasets, we recommend downloading the application from GitHub (https://github.com/sreejatadutta/PerSEveML) and running it locally on your system.

